# High-Resolution Genome-Wide Maps Reveal Widespread Presence of Torsional Insulation

**DOI:** 10.1101/2024.10.11.617876

**Authors:** Porter M. Hall, Lauren A. Mayse, Lu Bai, Marcus B. Smolka, B. Franklin Pugh, Michelle D. Wang

## Abstract

Torsional stress in chromatin plays a fundamental role in cellular functions, influencing key processes such as transcription, replication, and chromatin organization. Transcription and other processes may generate and be regulated by torsional stress. In the genome, the interplay of these processes creates complicated patterns of both positive (+) and negative (-) torsion. However, a challenge in generating an accurate torsion map is determining the zero-torsion baseline signal, which is conflated with chromatin accessibility. Here, we introduce a high-resolution method based on the intercalator trimethylpsoralen (TMP) to address this challenge. We describe a method to establish the zero-torsion baseline while preserving the chromatin state of the genome of *S. cerevisiae*. This approach enables both high-resolution mapping of accessibility and torsional stress in chromatin in the cell. Our analysis shows transcription-generated torsional domains consistent with the twin-supercoiled-domain model of transcription and suggests a role for torsional stress in recruiting topoisomerases and in regulating 3D genome architecture via cohesin. Significantly, we reveal that insulator sequence-specific transcription factors decouple torsion between divergent promoters, whereas torsion spreads between divergent promoters lacking these factors, suggesting that torsion serves as a regulatory mechanism in these regions. Although insulators are known to decouple gene expression, our finding provides a physical explanation of how such decoupling may occur. This new method provides a potential path forward for using TMP to measure torsional stress in the genome without the confounding contribution of accessibility in chromatin.

## Introduction

The double-stranded helical structure of DNA inherently results in the generation of torsional stress during a host of basic processes, including replication and transcription. Many essential motor proteins, such as RNA polymerase (RNAP), track the DNA helical groove during translocation, producing a relative rotation between the motor and the DNA. If RNAP rotation is restricted during transcription, DNA itself must twist, unwinding DNA to generate (-) supercoiling behind the polymerase, while overwinding DNA to generate (+) supercoiling in front, as described by the twin-supercoiled domain model of transcription^1–6^. Excessive torsional stress generated during transcription may, in turn, present challenges to motor progression^2,3,5–7^. Topoisomerases, a class of enzymes that can relax torsional stress and alter DNA topology, are essential to ensure cell cycle progression and are required for efficient transcription to occur^8–14^. However, residual supercoiling may offer advantages to cell survival. For example, (-) torsional stress behind these rotary motors may slow motor progression^2,3,14^ while promoting processes that require DNA unwinding, such as transcription initiation, replication, or DNA repair^6,15–20^. (+) torsional stress in front of these rotary motors, on the other hand, may slow motor progress^2,3,14,21^, dissociate proteins ahead of advancing motors^22^ and promote genome organization^23–25^.

Importantly, torsion can act over a distance so that torsion generated from one gene can regulate the transcription of a neighboring gene. For divergent genes, (-) torsion generated from the transcription of one gene can facilitate DNA unwinding of the promoter of the neighboring gene and thus promote its expression^5,6,26^. Interestingly, some promoter architectural factors, including sequence-specific transcription factors (ssTFs), can decouple the transcription between neighboring genes, effectively acting as insulators^27,28^. Although the precise mechanism of insulation remains unclear, it raises the possibility that ssTFs can insulate torsion between neighboring genes. Addressing such questions requires sensitive techniques to investigate torsional stress across the genome *in vivo*.

Elucidating the impact of torsional stress in cells requires accurate mapping of torsion. This requires a sensor that ideally should interact with the DNA to report its torsional stress, have fast and tunable binding kinetics for capturing a snapshot of the cellular state, be readily delivered into the cell, and have a robust readout. These requirements are nearly all met by 4,5′,8-trimethylpsoralen (TMP), a small photoreactive DNA intercalator that can cross-link to DNA upon UVA irradiation, with the extent of cross-linking modulated by TMP concentration and UVA dosage^29^. Because TMP unwinds DNA upon intercalation, its binding affinity, and thus the probability of generating interstrand crosslinks (ICLs) at a given site, is influenced by local torsional stress. Previous studies suggest that the relationship between torsion and the crosslinking rate is approximately linear across a relevant range of torsional stress^30,31^. While psoralen shows great promise for mapping torsional stress, significant challenges remain to be addressed before its full utilization can be realized.

Previous studies have laid the foundation for using TMP and other psoralen derivatives as torsional sensors^30,32–35^. However, factors besides the torsional state influence the resulting photo-binding distribution^30^. Importantly, the torsion signal mapped using TMP contains a zero-torsion baseline that varies across the genome, and this baseline is dependent on DNA sequence and TMP accessibility. Establishing this baseline is crucial for differentiating between (-) torsion and (+) torsion states across the genome, and in the absence of the zero-torsion baseline, the contribution from torsion to TMP intercalation may be conflated with TMP accessibility^36^. However, it is challenging to establish such a baseline as experiments must be performed under conditions that preserve both the chromatin state and the binding states of other proteins. Thus, interpreting the psoralen sensor readout requires a method to establish the zero-torsion distribution, but such methods have yet to be firmly established.

The impact of underlying DNA sequence on the zero-torsion psoralen binding landscape has been previously addressed by comparing the results from psoralen photo-binding in the cell to that of purified genomic DNA^30,32,33,35,36^. While this method corrects for the intercalation sequence-dependence, it does not account for the reduction of photo-binding due to restricted accessibility caused by bound proteins or chromatin. Indeed, previous studies have shown that chromatin reduces global psoralen crosslinking, suggesting that local effects of accessibility may be even more pronounced^30^. One method to assess the zero-torsion baseline is to digest the DNA using a DNA-nicking enzyme^34^. Still, this approach cannot ensure the preservation of the DNA or chromatin state before psoralen photo-binding. Other methods modulate RNAP or topoisomerase activities to generate a comparative map of torsional stress to extrapolate what might be a zero-torsion state but do not solve the problem at its core^32–35^. Importantly, alteration of normal cellular functioning can, in turn, alter DNA accessibility or the zero-torsion state. These complications give a clear motive for devising a strategy of measuring the zero-torsion baseline of TMP to reveal a more accurate signal of torsional stress.

To directly address this limitation, we report a method that enables psoralen mapping on relaxed DNA *in situ* while preserving the chromatin state at high resolution. We demonstrate the application of this method using *S. cerevisiae,* but the method is generalizable to other organisms. We found that the zero-torsion baseline contains significant contributions from individual nucleosomes within genes. After removing this baseline, we obtained a torsion signal that identifies regions of the genome under (-) torsion or (+) torsion. This revealed the twin-supercoiled domain model of transcription and allowed examination of how torsion may facilitate the recruitment of topoisomerases. We also show direct evidence for the presence of (+) torsion at cohesin-mediated loop boundaries, implicating its role in loop anchoring. Finally, we show evidence that sequence-specific transcription factor (ssTF) insulators prevent crosstalk between divergent promoters, providing a possible explanation for transcriptional coupling between promoters that lack bound insulators. This method provides a potential path forward for measuring torsional stress using psoralen-based methods in any cellular context.

## Results

### A method to establish the zero-torsion baseline

To establish the zero-torsion baseline for the psoralen signal, we must establish a method to account for psoralen’s sequence preference and accessibility without torsion under conditions that preserve the native-bound protein landscape (Fig. Supplement 1). To do this, we developed a method to prepare the chromatin zero-torsion state *in situ* and applied psoralen photo-binding (Fig. 1a, right panel). Formaldehyde preserved the chromatin landscape and transiently bound proteins (Materials and Methods) while stopping any subsequent cellular processes. The fixed cells were permeabilized so enzymes could diffuse into the nucleus. For *S. cerevisiae*, the cell wall was gently digested to create spheroplasts. To release torsion in intact spheroplast nuclei, we utilized DpnII, chosen for its high cut site density but inability to processively digest DNA, thus maintaining psoralen’s crosslinking ability. Upon DNA digestion, we observed an average fragment size of ∼2 kb, sufficiently short to release torsion but long enough to provide a chromatin substrate for psoralen binding. This zero-torsion state preparation was subsequently crosslinked using 365-nm light, enabling measurement of the torsion-free TMP intercalation landscape (referred to as TMP signal without torsion). The *in vivo* measurement (TMP signal with torsion) was performed by directly adding TMP to the yeast media since unmodified TMP is cell-wall permeable. Cell permeability enables photo-crosslinking to be performed before cells are harvested, thus capturing native cell activity (Fig. 1a, left panel).

**Figure 1.**
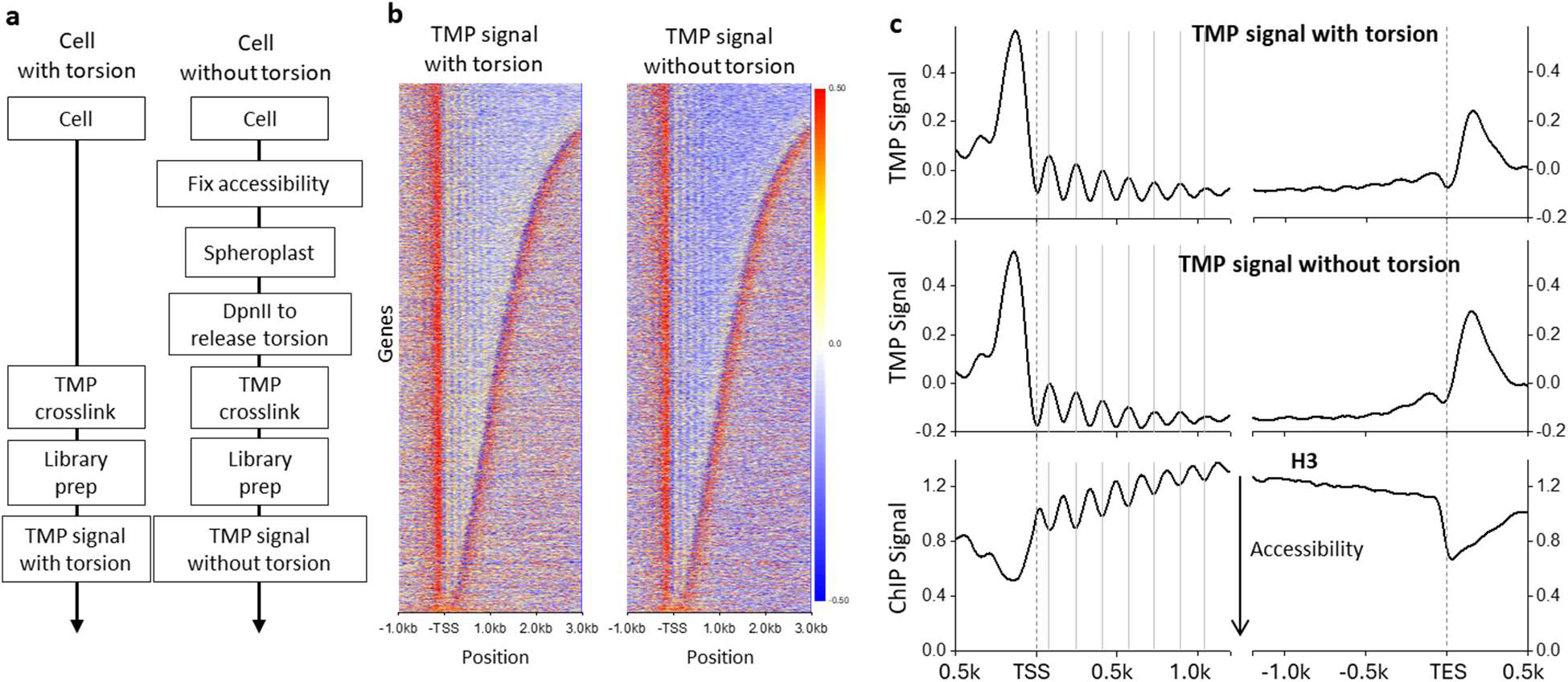
Measurement of the zero-torsion TMP baseline. (a) Strategy for determination of the TMP signal with and without torsion. (left) Cells with torsion are crosslinked with TMP prior to library preparation for the TMP signal measurement. (right) To measure the TMP baseline signal without torsion, the native-bound protein landscape is fixed by formaldehyde, and the torsion is released via DpnII digestion of the spheroplast cells before TMP crosslinking. Both TMP crosslinking samples are then mapped using Illumina-based sequencing to generate the TMP signal (Materials and Methods). (b) Heatmaps of TMP signals aligned at the TSSs of 5,925 genes for both the TMP signal with torsion (left, *n* = 2 biological replicates) and the TMP signal without torsion (right, *n* = 3 biological replicates) with rows sorted by the ORF size. The TMP signal was normalized against that of the purified DNA. (c) (top) Average TMP signals with and without torsion from data in (b) aligned at the TSS (left) and aligned at the TES (right). For comparison, previous H3-ChIP-MNase-Seq data^27^ are also shown (bottom). To minimize signal bias from neighboring genes, signals downstream of the TES were excluded from the analysis of the left panel plots, and signals upstream of the TTS were excluded from the analysis of the right panel plots.

To locate psoralen crosslinks on the DNA, we utilized an exonuclease-based enrichment strategy followed by an Illumina-compatible library prep, similar to TMP-seq^30,33^. Exonuclease-based enrichment relies on interstrand crosslink (ICL) formation, which prevents DNA denaturation before digestion by the single-strand-specific ExoI. We use the digested fraction to estimate the crosslinking density and provide quality control for ICL enrichment over the background. As controls, we found that libraries prepared without psoralen photo-binding showed a very low signal level (Fig. Supplement 2). By tuning the UV dosage, we ensured that the average densities of the *in vivo* and zero-torsion measurements were comparable at ∼0.1-0.25 ICLs/kb (Fig. Supplement 2). If ICL density is too low, the signal-to-noise ratio is poor; if ICL density is too high, there can be library preparation artifacts (Fig. Supplement 3). ICL densities within the range used display remarkably similar signals (Fig. Supplement 2). This allows the TMP signals with and without torsion to be compared to obtain an accurate genomic map of psoralen binding due to torsional stress. In addition, we conducted a control experiment using purified DNA to assess the extent of psoralen crosslinking due to DNA sequence preference (Fig. Supplement 2). This TMP signal was only used during an intermediate analysis step to isolate the psoralen intercalation preference not due to DNA sequence (Fig. 1b,c).

### TMP allows the mapping of positioned nucleosomes

To evaluate the contributions of DNA accessibility to the TMP intercalation profile, we examined the TMP signal downstream of transcription start sites (TSSs), which are known to have positioned nucleosomes^37,38^. We expect TMP to preferentially intercalate in linker DNA regions between nucleosomes in samples with and without torsion, and the ability to reveal these well-positioned nucleosomes is a measure of our TMP data quality. We found a striking similarity between the signals from the two samples (Fig. 1b). This finding demonstrated that the *in vivo* psoralen signal is likely dominated by accessibility so that TMP intercalation preference due to torsion may represent a smaller contribution to the overall signal than protein-mediated DNA accessibility. Therefore, using the TMP intercalation signal alone to indicate torsion may be prone to interpretation artifacts.

For both datasets, we detected a TMP peak upstream of the TSSs, with the peak amplitude being slightly larger for the condition with torsion. This peak coincides with the nucleosome-free regions (NFRs), which should be more accessible to TMP (Fig. 1c). Within the gene bodies, we also observed a clear ∼160 bp periodic signal, similar to that of the positioned nucleosomes near the promoters (Fig. 1c). However, the periodic pattern anti-correlates with H3-ChIP-MNase seq data, indicating that psoralen prefers to bind in the linker regions between nucleosomes or in NFRs (Fig. 1c)^27^. The decay in the amplitude of the somewhat sinusoidal TMP signal away from the TSS mirrors the decay in the amplitude of the H3-ChIP-seq signal, reflecting a decrease in nucleosome phasing further from the promoters^38^. Immediately downstream of the transcription end sites (TESs), we observed a smaller TMP peak than before the TSSs, likely due to the presence of promoter regions associated with a downstream codirectional gene. These findings show that the TMP intercalation distribution reflects many features previously observed using other chromatin accessibility assays and suggest that the TMP signal may be dominated by chromatin accessibility.

### Twin-supercoiled domains of transcription

To determine psoralen intercalation due to DNA torsion, we subtracted the zero-torsion baseline from the *in vivo* distribution, eliminating contributions from sequence preference and accessibility (Fig. 2a,b). The resulting signal has a ∼ 5-fold reduced peak-to-valley amplitude and should reflect contributions solely due to torsional stress. Most of the nucleosome signal was abolished within the gene bodies, indicating that most accessibility bias has been removed, although a small residual remained (Fig. 2b). We refer to the corrected signal as the “torsion signal,” with a positive value indicating (-) torsion, which facilitates psoralen intercalation, and a negative value indicating (+) torsion, which restricts psoralen intercalation. Using this approach, we examined whether we could detect the torsional state more accurately during transcription. To examine the torsion distribution along a gene, we focused on regions near promoters and terminators. We found the torsion signal now reveals (-) torsion upstream of promoters and (+) torsion downstream of terminators (Fig. 2b). This may be interpreted as a statistical average of a bulk population of Pol II molecules, which have elongated to random locations along the gene in different cells. Since each Pol II molecule generates (-) torsion behind and (+) torsion ahead, according to the twin-supercoiled-domain model, the net torsion signal from different Pol II molecules averages to near zero, except at gene boundaries. We also found that torsion is confined to within 1-2 kbp of the promoters and terminators, supporting previous findings that transcription-dependent dynamic supercoiling is a short-range genomic force^32^.

**Figure 2.**
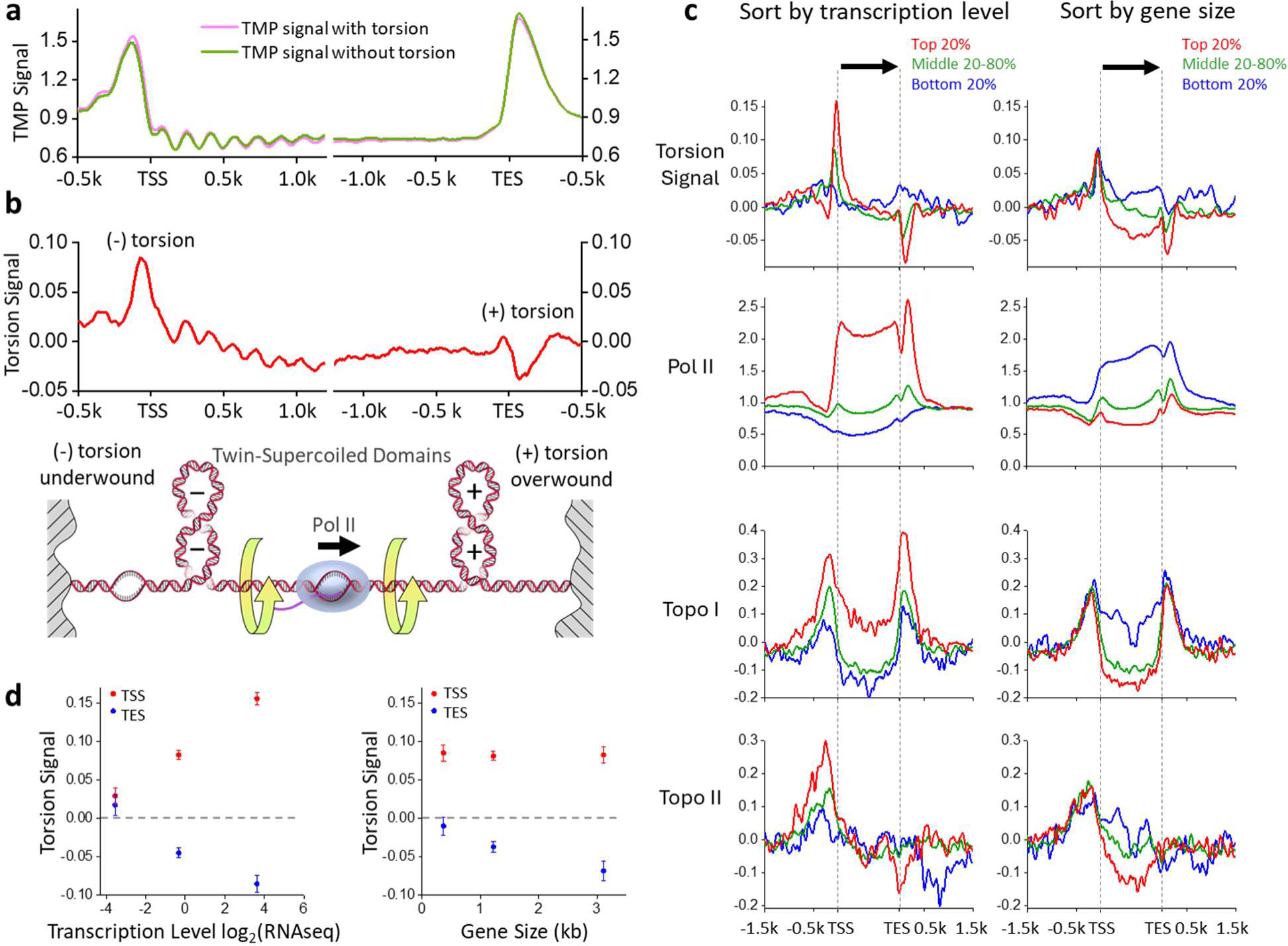
The twin-supercoiled domains of transcription. (a) Average TMP signals with and without torsion using data from Fig. 1c, replotted for easy comparison, except without normalization against that of the purified DNA. (b) The torsion signal, defined as the difference between the TMP signal with torsion and the TMP signal without torsion. The bottom cartoon shows the twin-supercoiled domain model of transcription. (c) The torsion signal of genes sorted by transcription level and gene size, with genes grouped by percentiles of mRNA abundance and ORF size, respectively. Positions within the gene body are scaled by the gene length while preserving the torsion signal amplitude (Materials and Methods). The torsion signal is compared to previous data of Pol II (Rpb3-ChIP-Exo)^27^, topo I (Top1-ChIP-Exo)^27^, and topo II (Top2-ChIP Exo)^27^. (d) Mean value of the torsion signal averaged within 20 bp of either the peak near the TSS (red) or the valley near the TES (blue) as functions of the average gene expression level (left) or gene size (right) within the percentile groups specified in (c). Error bars are standard errors of the means.

To further examine whether the torsion signal is due to transcription, we classified genes into percentile groups based on their expression level (Fig. 2c,d). We expected that the average transcription-generated torsional stress should increase with increased gene expression. Thus, we sorted the genes based on their expression level according to previous mRNA-seq data^39^. Consistent with this prediction, genes in the bottom 20% of the expression level do not display a significant peak near the promoters nor a significant valley near the terminators. As the gene expression level increases, the torsion signal amplitudes near the promoters and terminators monotonically increase on average (Fig. 2c,d). As expected, gene expression strongly correlated with Pol II presence along the genes (based on previous ChIP-seq data of Pol II^27^). This shows that active transcription accumulates torsional stress near highly transcribed genes, supporting earlier findings from *in vitro* and *in vivo* studies^2,4–7,40–42^.

The persistence of torsion near genes, even with full native topoisomerases, indicates that topoisomerases cannot fully keep up with transcription activities. To examine the correlation of torsional stress with topoisomerase activity, we analyzed previous ChIP-seq data of topoisomerase I (topo I) and topoisomerase II (topo II)^27^, which are the primary enzymes for relaxing transcription-generated torsion. Topo I binds near promoters and terminators, and its enrichment level correlates with an increase in the torsion signal (Fig. 2c). In contrast, topo II binds near promoters but not near terminators. As the torsion signal increases, the topo I and topo II enrichment levels also increase, suggesting that torsion might facilitate topoisomerase recruitment or retention. Unlike topo I, which binds to one DNA segment, topo II relaxes DNA via a strand-passage mechanism^43^ and has been found to prefer binding to a DNA crossover^13,44–47^. Because the DNA in a nucleosome wraps around the histone octamer in a left-handed fashion, the nucleosome conformation exhibits a chiral response to DNA supercoiling^48,49^. While (-) torsion readily enables DNA crossing formation via the juxtaposition of the entry and exit DNA segments in a nucleosome, (+) torsion displaces entry and exit DNA segments before bringing them together again, buffering torsional stress^14,48^. Thus, there is less DNA crossing at a nucleosome available to allow topo II binding. Consistent with this interpretation, we previously found that topo II has a faster relaxation rate and is more processive on chromatin under (-) supercoiling than under (+) supercoiling^44^. Thus, the absence of topo II near the terminators may not be due to any active exclusion of topo II at those regions but is a natural consequence of the interplay between the DNA chirality and nucleosome chirality.

Previous studies suggest torsion accumulates more significantly over longer genes^5^. As Pol II elongates, its rotation may be more restricted with a lengthening in the RNA transcript, which is known to be associated with large machinery, such as spliceosomes^50^. Thus, torsional stress might significantly increase toward transcription termination for longer genes. We grouped data based on gene size (Fig. 2c,d), and consistent with this prediction, the (-) torsion near the promoters remained unchanged with an increase in gene size, but the (+) torsion near the terminators increased with gene size to a moderate extent. As expected, we found no significant dependence of topo I or topo II enrichment on gene size near the promoters. However, we noticed a similar lack of dependence near the terminators, likely due to the moderate dependence of torsion on gene size.

### Torsion accumulation between genes

Transcription-generated torsion may dissipate over distance due to topoisomerase relaxation and DNA end rotation^6,32,51^. If so, torsion generated from one gene will most likely impact the torsional state of neighboring genes. For example, (-) torsion may more likely accumulate near promoters of two divergent genes, while (+) torsion may more likely accumulate near terminators of two convergent genes. To examine this prediction, we plotted the torsion signal in three gene-pair configurations (Fig. 3). We mapped the average behavior of torsional stress in the intergenic regions after rescaling the intergenic distance between nearest neighbors. We found that between promoters of two divergent genes, the torsion signal rises throughout the intergenic region, consistent with (-) torsion accumulation between these gene pairs. Conversely, we observed that between terminators of converging genes, the torsion signal drops throughout the intergenic region, consistent with (+) torsion accumulation between these gene pairs. When two neighboring genes are oriented codirectionally, the torsion signal drops after the terminator of the trailing gene and rises before the promoter of the leading gene, consistent with generation of (+) torsion by the trailing gene followed by generation of (-) torsion by the leading gene. Thus, by grouping neighboring genes in these configurations, we demonstrate how torsion accumulates between neighboring gene pairs.

**Figure 3.**
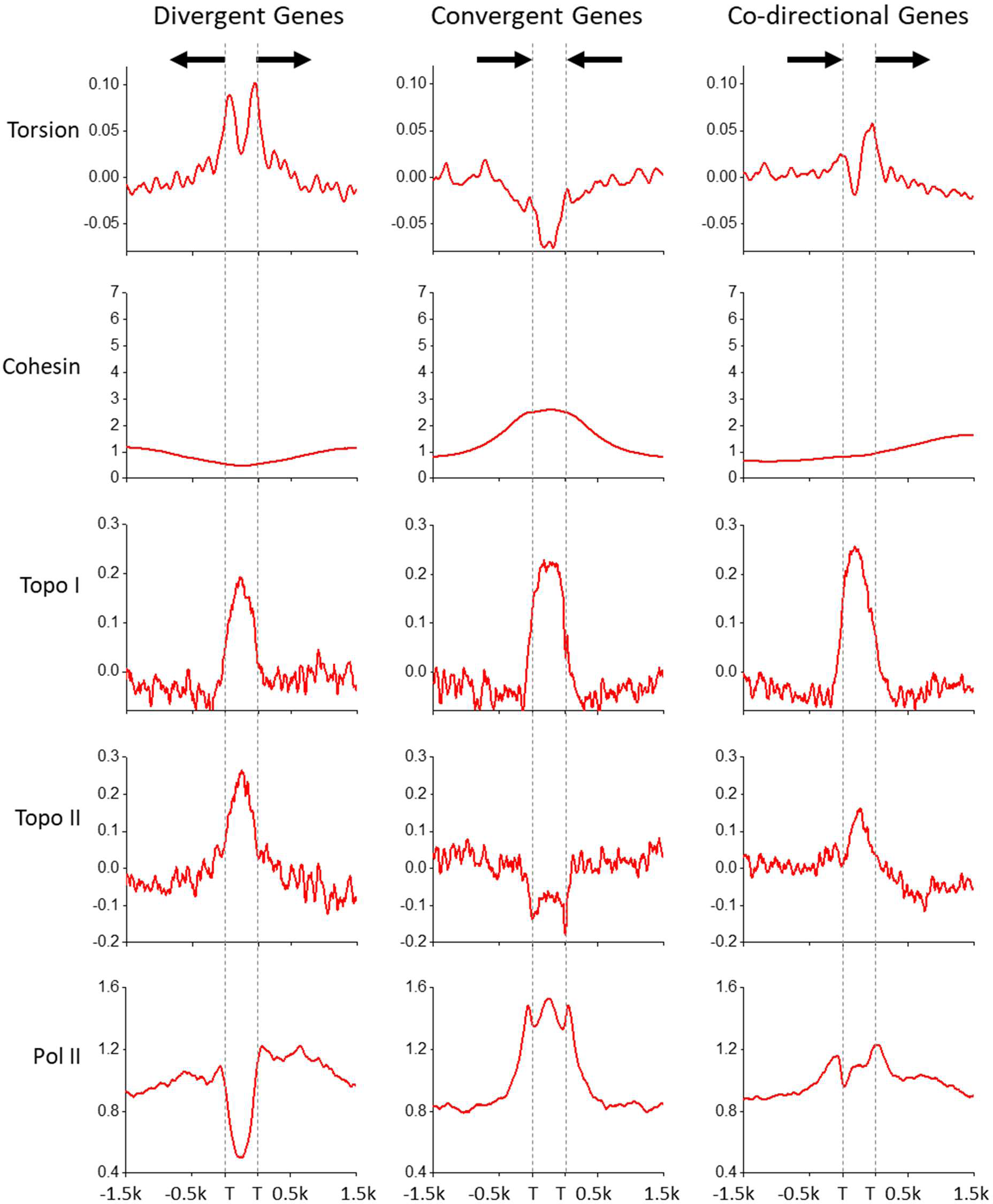
**Torsion between genes.** The torsion signal plotted for three gene-pair configurations: divergent (left, 1275 pairs), convergent (middle, 1364 pairs), co-directional (right, 2426 pairs). The torsion signal is compared to previous protein mapping data of the same gene-pair configurations for cohesin (SccI-Chip)^24^, topo I (Top1-ChIP-Exo)^27^, and topo II (Top2-ChIP Exo)^27^, and Pol II (Rpb3-ChIP-Exo)^27^. TSS or TES is indicated as “T”.

Since transcription-generated torsional stress hinders Pol II progression, topoisomerases must be recruited to active genes to resolve the torsional stress. To examine where Pol II, topo I, and topo II are located, we analyzed previous ChIP-seq data of Pol II, topo I, and topo II^27^. We show that Pol II has a minimal presence in the intergenic region between divergent genes and is enriched in the intergenic region between convergent genes, consistent with a previous finding that after termination, Pol II tends to remain on the DNA downstream of the terminator^27,39^.

We also find that topo I is enriched in all intergenic regions with significant (+) torsion or (-) torsion, indicating that topo I effectively targets regions under torsional stress. In contrast, topo II localizes to the intergenic region between divergent genes but is depleted from the intergenic region between convergent genes, indicating that topo II prefers regions with (-) supercoiling but is restricted from regions with (+) supercoiling, further supporting our interpretations above. These data further demonstrate how topo I and topo II may differentially target diverse environments in the genome.

### The role of torsion in genome structure

Maintenance of three-dimensional genome structure is a crucial step in cellular function. Recent yeast genome studies have found mitotic loops near the boundaries of convergently oriented gene pairs^23,24,52^. This suggests that (+) torsional stress may regulate genome folding. We examined the signal near loop boundaries (Fig. 4a) by aligning all loop boundaries and grouped signals based on the looping strength of previously published MicroC-XL Data^52^. This sorting method showed that cohesin^24^ is enriched at the two-loop boundaries as expected. Using this sorting method, we found (+) torsion signal at the loop boundaries, which also coincide with regions of convergent gene pairs. We found that (+) torsion correlates with strengthening cohesin loop boundaries, suggesting that (+) torsion in these regions may facilitate cohesin recruitment (Fig. 4a, cartoon). These data provide evidence that the (+) supercoiling generated by transcription may facilitate genome folding in coordination with other participating proteins.

**Figure 4.**
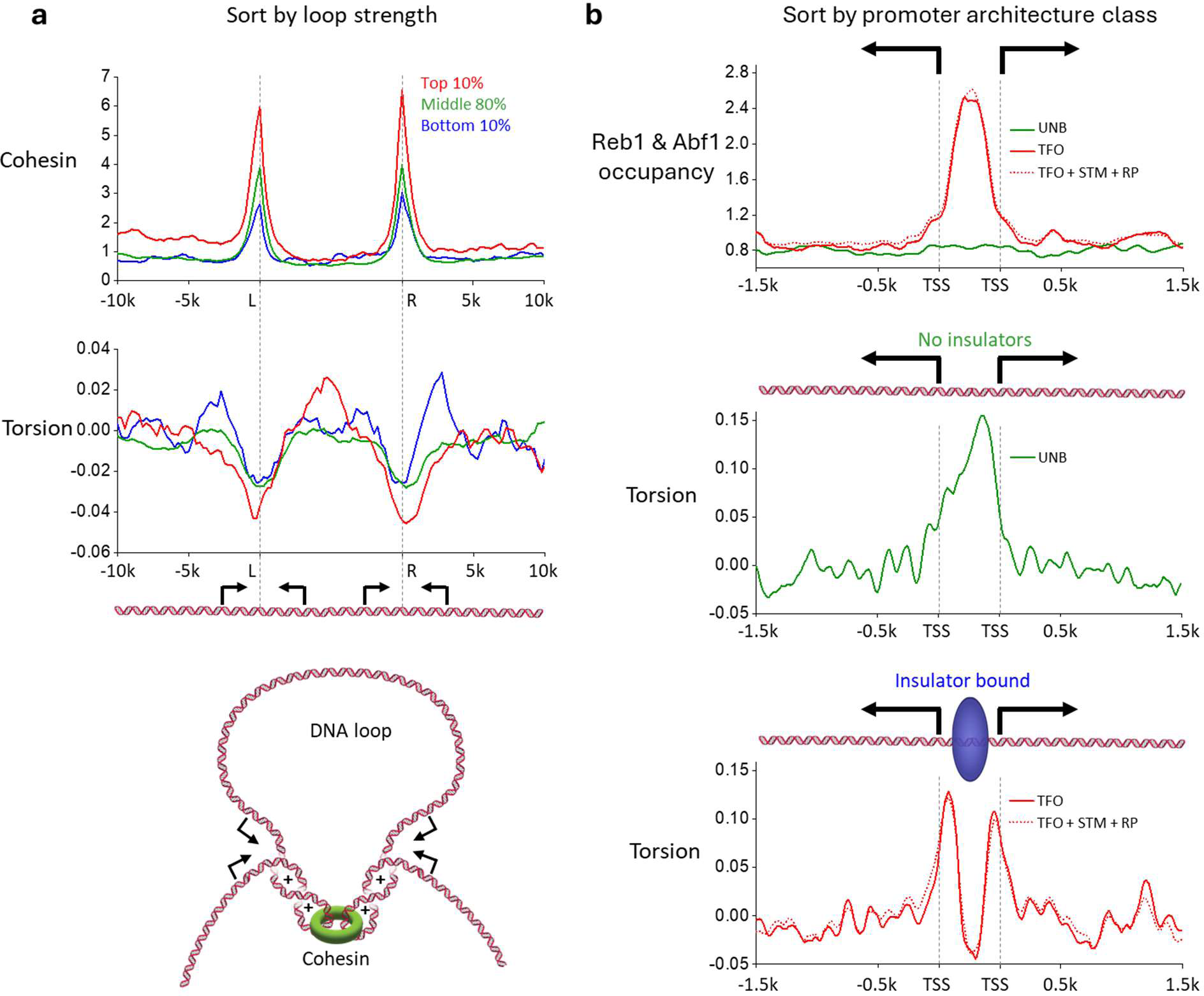
Torsion impacts 3D genome structure and couples gene expression. (a) Cohesin-ChIP seq^24^ and torsion signals plotted across 527 detected mitotic loops from previous MicroC-XL Data^52^, grouped by percentiles of the loop score. Positions between the two loop boundaries (L and R) are scaled by the DNA length between the boundaries while preserving the torsion signal amplitude (Materials and Methods). Below is a cartoon depicting a DNA loop anchored by cohesin at sites of (+) supercoiling. (b) Average ChIP-Exo signal of Abf1 and Reb1^27^ (top plot), and torsion plotted across subsets of divergent genes (middle and bottom plots). Gene pairs are separated by defined classes^27^ where both genes in the pair are of the same class as follows: unbound class (UNB; green, 421 pairs), ssTFs only (TFO; red-solid, 320 pairs), ssTFs and other factors (TFO, STM, and RP; red-dotted, 439 pairs). Positions between the two divergent promoters are scaled by DNA length between the two promoters while preserving the signal amplitude (Materials and Methods).

### Insulators decouple torsion between divergent genes

While a single (+) torsion intergenic region is present between two convergent genes, we detect two (-) torsion intergenic regions between two divergent genes (Fig. 3). The presence of the two (-) torsion peaks indicates that (-) torsion from these promoters is somewhat decoupled. However, it is unclear what limits the diffusion of (-) torsion between the two promoters. Thus, we systematically examined the promoter regions of different promoter architectures to identify factors that may play a role.

Most promoters in *S. cerevisiae* are not regulated by ssTFs and thus unbound by anything besides a PIC (termed “UNB”)^27,53^. Interestingly, our data show a single elevated (-) torsion region between this UNB class of divergent promoters (Fig 4b, upper and middle plots), suggesting that torsion generated from one gene can impact the expression of its neighboring gene, consistent with previous findings that the expression of these genes is coupled^27,53^.

Another large class of *S. cerevisiae* promoters (termed “TFO”) are regulated by insulator ssTFs, such as Reb1 and Abf1, which decouple interactions between neighboring genes^27,28,53–55^. A minority of genes, such as inducible promoters (termed “STM”), are even more highly regulated by a host of additional transcription factors and usually contain an upstream activating sequence^27^. Strikingly, these bound promoters show two distinct (-) torsion peaks, suggesting that the insulators can insulate torsion between the two genes (Fig 4b, upper and lower plots). This result indicates that bound insulators may be anchored to cellular structures in a manner that prevents rotation of the bound DNA segments and thus blocks the progression of torsion from one promoter to the next. Given that DNA unwinding facilitates transcription initiation, insulators may prevent torsional crosstalk between adjacent divergent promoters while also preventing TATA-binding protein (TBP) from sliding back and forth between promoters.

## Discussion

This work developed a method to profile the *in situ* zero-torsion baseline of psoralen-based mapping of torsional stress in yeast. The approach enabled high-resolution mapping of the psoralen intercalation distribution on torsionally relaxed DNA, while preserving the protein-bound state of chromatin. This preservation is crucial to correcting the psoralen-based torsion map because it allows the removal of the contributions of both chromatin accessibility and DNA sequence. Through this process, we found that the psoralen technique is excellent at generating high-resolution accessibility maps of chromatin, resolving positioned nucleosomes at a resolution superior to that of ATAC seq^56^. This demonstrates the potential utility of psoralen well beyond torsional measurements.

Our resulting torsion signal also reveals a characteristic torsion distribution of the twin-supercoiled-domain model of transcription with (-) torsion behind promoters and (+) torsion in front of terminators. We demonstrate how torsion increases with gene expression, and topoisomerases are recruited to regions with torsion. We also provide an explanation for the differential enrichment of topo I and topo II based on the torsional mechanics of chromatin. In addition, we show that cohesin loop boundaries coincide with regions of (+) torsion in the genome, indicating a role of torsion in helping cohesin restructure the genome topology.

Importantly, we provide a possible physical explanation for how ssTF insulators may decouple highly regulated divergent genes and insulate gene expression. We show that (-) torsion diffuses across promoter regions lacking ssTFs, while (-) torsion is restricted to their respective promoters in the presence of ssTFs. These findings suggest that these ssTFs may insulate torsion between the promoters to decouple their expression.

Accessibility is likely to greatly influence the readout of any torsion sensor that relies on DNA binding. Thus, establishing the zero-torsion baseline is an essential step and must be performed under a condition that keeps chromatin intact. While torsion sensors other than TMP have been applied to generate genomic maps, these sensors, such as biotin-functionalized psoralen derivatives^34–36^ and the bacterial protein GapR^57^, are larger in size than TMP, and thus, these sensors are likely also sensitive to DNA accessibility. Since the method we developed to obtain the zero-torsion state is not specific to any sensor, it may be applied to other torsion sensors. Additionally, although this study utilized budding yeast as a model organism, preparation of the zero-torsion state can be similarly applied to cells of other organisms. Our work shows a way to measure torsion more accurately and helps to lay a foundation for future quantitative studies of torsion in the genome.

## Materials and Methods

### Cell Culture

The *Saccharomyces cerevisiae* strain BY4741 (*MAT a his3Δ1 leu2Δ0 met15Δ0 ura3Δ0 bar1::NatMX* his3Δ1::HIS3-ADH1pr-hENT-GPDpr-HSV-TK)^58^ was used for all experiments. Cells were grown at 30°C in YPD media until the OD600 reached ∼0.6, and then processed for different conditions as described below.

### Zero torsion state preparation

Cells designated as zero torsion state controls were crosslinked with 1% (v/v) formaldehyde (Fisher, F79-500) for 15 minutes at 30°C. Formaldehyde was quenched with 0.4 M glycine (Sigma, 357002) for 5 minutes at room temperature. Following quenching, cells were pelleted and washed 2X with PBS at 4°C. Next, cells were spheroplasted via 0.01% (v/v) 10 mg/µl zymolyase (Sunrise Science Products, N0766391) treatment in 1 M sorbitol (Sigma, S6021), 0.5 mM 2-mercaptoethanol (Sigma, 444203) for 20 minutes. Spheroplasts were confirmed to be >90% of cells using microscopy. Formaldehyde-fixed spheroplasts were centrifuged at 1,000 g for 10 minutes at 4°C, washed with 25 ml of cold PBS (Thermo, AM9624), and resuspended in 100 µl of 1x DpnII reaction buffer (NEB, R0543) containing 100 units of DpnII (NEB, R0543) and supplemented with 0.1% (v/v) NP-40 (Thermo, 85124), and 10 µM TMP (Sigma, T6137). Samples were incubated for 2 hours at 37°C followed by TMP photobinding as described below.

### Trimethylpsoralen Photobinding

The 4,5′, 8-trimethylpsoralen (TMP) was dissolved in 100% ethanol at a final stock solution of 2 mM. TMP was added to growing cells at a final concentration of 10 µM and cells were further incubated at 30°C for 1 hour. Twenty-five mL of cell solution was placed in a 10 cm petri dish and exposed to ∼0.15 W/cm^2^ of 365-nm light for 20 seconds. Directly after TMP photobinding, the cells were fixed in 95 ml 70% −20 °C ethanol and placed at −20°C overnight.

Cells were then harvested by centrifugation at 10,000 g for 10 minutes, resuspended in 500 µl of 50 mM sodium citrate (Sigma, 567446), pH 7.4, and incubated at room temp for 10 minutes. Centrifugation and wash steps were repeated once more, followed by a 20-minute Zymolyase treatment as detailed above. Spheroplasts were harvested by centrifugation and resuspended in 100 µl of lysis buffer (300 mM NaCl (Fisher, BP358), 50 mM EDTA (Fisher, AM9260G), 2% SDS (Sigma, 71725), and 5 % (v/v) RNase A (Thermo, 12091021) for ∼2 hours and then vortexed for 2 minutes. 10 µl of Proteinase K (NEB, P8107S) was added, and cells were incubated for 2 hours at 65 °C.

Zero torsion samples followed a similar photobinding protocol but were exposed to UV light at ∼0.35 W/cm^2^ for 10 seconds in a 1.5 ml Eppendorf tube. Samples were harvested at 10,000 g at 4°C, the supernatant was removed, and they were resuspended in 100 µl of Lysis buffer containing 10 µl Proteinase K. The zero torsion samples were incubated at 65°C overnight to reverse formaldehyde crosslinks. After lysis, genomic DNA was extracted from all samples via 2X phenol-chloroform (Sigma, 77617) extraction followed by isopropanol (Sigma, 59304) precipitation.

### TMP-seq

The TMP-seq protocol was adapted from previous work^30,33^. Extracted genomic DNA was sonicated to ∼400 bp fragments using appropriate settings on a Covaris E200 sonicator. Following sonication, DNA was treated with 5 µl RNase A and purified using a Zymoclean DNA Clean and Concentrator-5 kit according to the manufacturer’s protocol using a 7:1 binding buffer:sample ratio. After quantification using a Biotum HS DNA quantification kit, 250 ng of DNA was diluted into 86 µl of 10 mM Tris-Cl pH 8.5, incubated at 98°C for 15 minutes, and then directly moved to ice for 2 minutes. DNA was then digested with 4 µl of ExoI (NEB, M0293) in 1X ExoI reaction buffer at 37°C for 1 hour. This digestion process was repeated by incubating at 98°C for 1 hour and then adding an additional 4 µl of ExoI followed by another incubation at 37°C for 1 hour. Digested samples were purified using a Zymo DNA Clean and Concentrator-5 kit using a 7:1 binding buffer: sample ratio. DNA was again quantified using the Biotum HS DNA assay. Illumina adaptors were ligated to the digested samples using the NEBNext Ultra II DNA library prep kit for Illumina (NEB, E7645) at half the standard total reaction volume of the manufacturer’s protocol. Adapter ligated DNA was purified with 2X Ampure beads (Beckman Coulter) and eluted in 40 µl of 10 mM Tris-Cl pH 8.5. DNA was stored at −20 C until further processing.

To locate psoralen interstrand crosslinks, DNA from the previous step was incubated with 5 µl of lambda exonuclease (NEB, M0262) in the provided buffer in a 50 µl total reaction volume at 37°C for 1 hour, purified with 1.8X Ampure beads, and eluted with 30 µl of 10 mM Tris pH 8.5. Lambda-digested DNA was then subjected to 10 rounds of primer extension with Q5 DNA polymerase (NEB, M0491), using 10 nM of the primer (ID): 5’-GTGACTGGAGTTCAGACGTGTGCTCTTCCGATC*T on a thermocycler (98°C for 10s → 10 rounds (98°C, 10s → 65°C, 2 min)→ 65°C, 5min → 95°C 1min → 4°C). Products were purified using 1.8x Ampure beads and eluted in 40 µl 10mM Tris pH 8.5. Primer extension products were ribotailed using 1.4 µl Terminal Transferase (NEB, M0315) and 0.4 mM rGTP (Roche, 11277057001) in Terminal Transferase buffer supplemented with 0.25 mM CoCl_2_ by incubating at 37°C for 30 minutes followed by incubation at 75°C for 20 min. Ribotailed products were purified with 1.8X Ampure beads, eluted with 30 µl 10mM Tris pH 8.5. Purified ribotailed products were ligated with 0.33 µM annealed adapter oligos (IDT) (Top: 5’p-AGA TCG GAA GAG CGT CGT GTA GGG AAA GAG TGT-3’, Bottom: 5’ ACA CTC TTT CCC TAC ACG ACG CTC TTC CGA TCT CCC −3’) using 1.2 µl T4 DNA ligase (NEB, M0202S) in the provided buffer for 1 hour at 16 °C, and subsequently purified with 1.8X Ampure beads and eluted in 20 µl 10mM Tris pH 8.5. Finally, products were indexed and amplified using NEBNEXT Multiplex Oligos for Illumina set 1 (NEB, E7600) following the manufacturers protocol with a total of 12 PCR cycles. The PCR products were purified using 0.8X Ampure beads and eluted in TE (Invitrogen). Samples were pooled at equimolar concentrations and empty adapters were removed from the pool.

### Sequencing and Data Processing

Pooled libraries were sequenced on a NextSeq 2000 (Illumina) using a NextSeq 2K P2 100 bp kit (Illumina) in paired-end mode with a 63 bp Read_1 and 59 bp Read_2. Sequencing, demultiplexing and Fastq generation were performed by the Cornell Biotechnology Resource Center. Fastq files were aligned to the SacCer3 Genome using BWA-mem2^59^ with default settings for paired-end reads. Sam files were converted to BAM files followed by fixmate and duplicate marking using Samtools^60^. Duplicates were again marked using picard-MarkDuplicates^61^, and duplicates were removed using Samtools-view. Bedgraph files were generated using deeptools^62^ bamCoverage with 10 bp bins using the following settings (--samFlagInclude 64--Offset 1) yielding the coverage of the 5’ end of Read_1 where the ICL was located.

Bedgraph files were further processed in MATLAB to ensure one bin per 10 bp interval, and normalized to the mean number of reads per bin. Replicate bedgraphs were averaged, and indicated subtractions were done using MATLAB. Normalized bedgraphs were converted to bigwig files using ucsc-bedGraphToBigWig^63^. Heatmaps and composite plots were generated using deeptools computeMatrix, and visualized using Origin Lab. Bed files containing coordinates of verified *s. cerevisiae* ORFs were downloaded from the Saccharomyces Genome Database (SGD)^64,65^, and a region blacklist was downloaded from the yeastepigenome.org^27^. Bed files for each gene configuration in Figure 3 were generated by considering the boundaries of nearest neighbor verified ORFs and classifying them based on their strand configuration. Regions that had any annotations according to the list of all yeast genes from the SGD were removed to isolate true intergenic regions. The resulting Bed files were used to make heatmaps and composite plots as described above.

### Data Processing

BAM files of ChIP-Exo, and H3-ChIP-MNase data was downloaded from yeastepigenome.org^27^. For ChIP exo, bigwig files were calculated using the 5’ end of read 1 as in^27^. Signals were then normalized to the mean number of reads per 10bp bin. For H3-ChIP-MNase, the center of the fragment was considered. Top1 and Top2 ChIP-Exo data were further processed by subtracting the No-TAP control (ChIP-Exo background). Previous Scc1 ChIP-seq IP and Input DNA data was processed by downloading bigwig files from GSE235560^24^. Deeptools bigwigCompare was used to normalize Scc1 ChIP/Input DNA.

mRNA-seq data was downloaded from GSE76142^39^ in bigwig format. The minus strand signal was subtracted from the plus strand signal. Expression level for a gene was calculated using mean absolute value of the subtracted signal within the ORF, bed files containing expression levels were used to separate data into percentile groups.

To determine positions of intrachromosomal loop boundaries and their scores, mcool files of MicroC-XL data were downloaded from GSE151553^52^. Boundaries were located and scored using cooltools dots^66^ with a 500 bp bin size. Loop calls were refined by only considering the smallest loops in nested configurations, thereby only representing each region once. Boundary positions were adjusted by locating the position of the nearest (<500 bp) Scc1-ChIP-seq peak on either loop boundary.

## Data and Material Availability

Raw and processed sequencing data files have been deposited to GEO under accession GSE284352. The custom code used to process the raw data and generate plots is available on GitHub (https://github.com/WangLabCornell/Hall_et_al).

## Acknowledgments

We thank members of the Wang Laboratory for helpful discussion and comments and the Cornell Biotechnology Resource Center for sequencing support. This work is supported by the National Institutes of Health grants R01GM136894 (to M.D.W.), T32GM008267 (to M.D.W.), and R35GM145217 (to B.F.P.). M.D.W. is a Howard Hughes Medical Institute investigator.

## Additional Information

### Contributions

Conceptualization, P.M.H and M.D.W; Methodology, P.M.H, L.B, M.B.S, B.F.P, and M.D.W; Software, P.M.H and L.A.M; Validation, P.M.H, L.A.M; Formal Analysis, P.M.H, L.A.M, and M.D.W; Investigation, P.M.H and L.A.M; Resources, L.B, M.B.S, and B.F.P; Data Curation, P.M.H and L.A.M; Writing – Original Draft, P.M.H and M.D.W; Writing – Review & Editing, P.M.H, L.A.M, B.F.P and M.D.W; Visualization, P.M.H. and M.D.W.; Supervision, M.D.W.; Project Administration, M.D.W.; Funding Acquisition, M.D.W.

---

**Figure Supplement 1.**
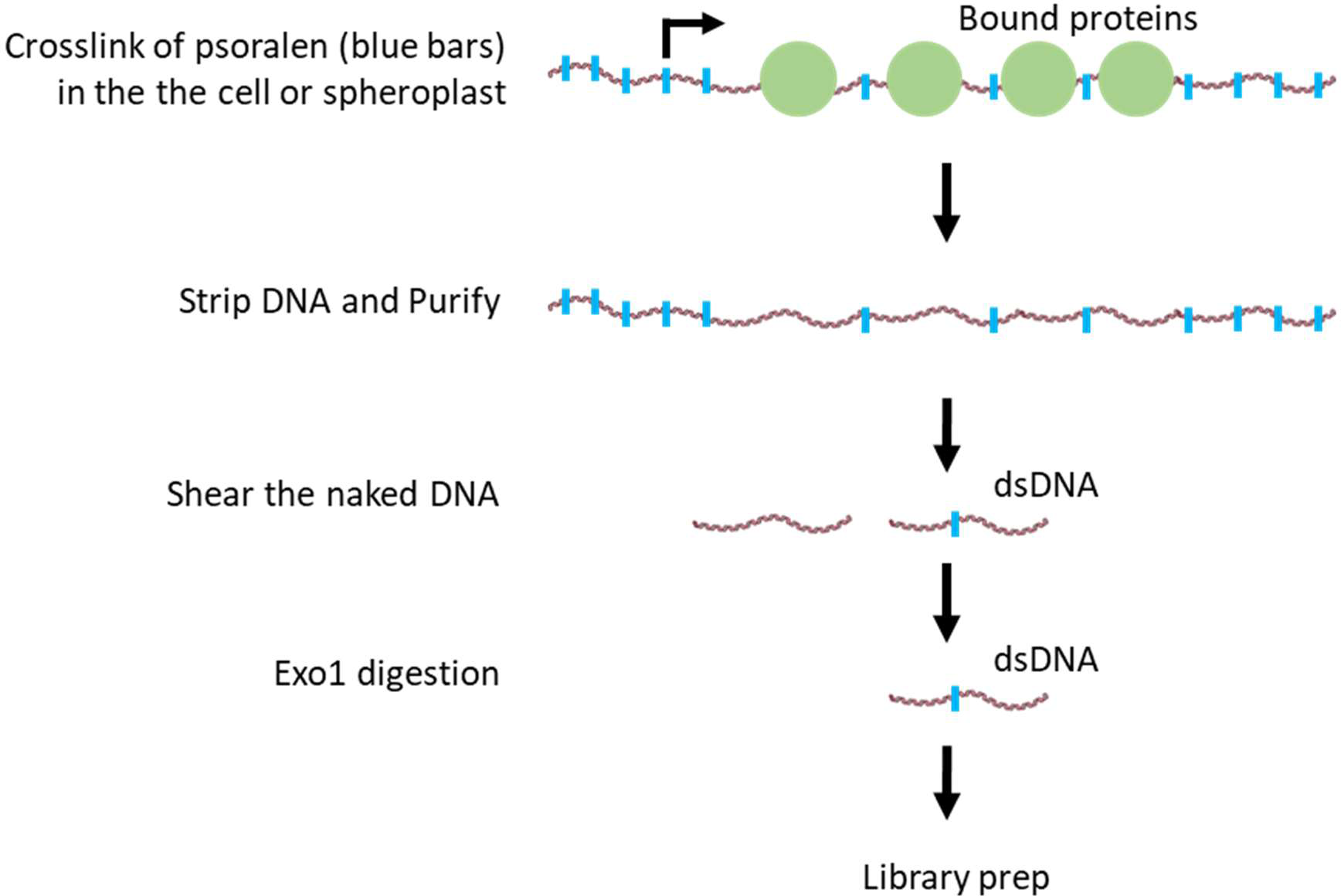
Psoralen binding and accessibility. Cartoon depicting the experimental method for mapping TMP cross-linking on genomic DNA. The TMP has preferential binding to naked DNA where there are no bound proteins, such as in nucleosome-free regions and on linker DNA.

**Figure Supplement 2.**
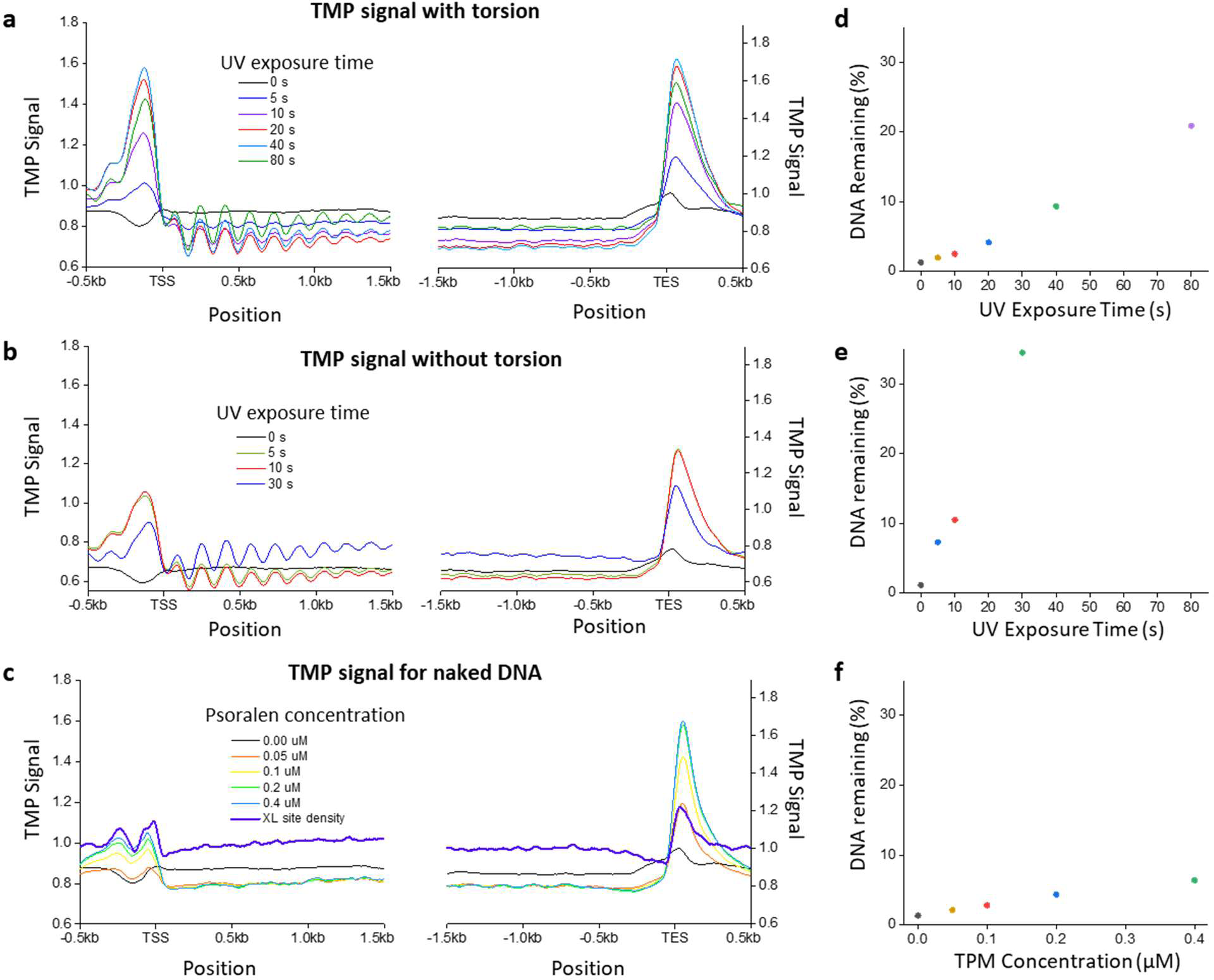
Psoralen Crosslinking Conditions. (a) Average TMP signal with torsion of all 5925 genes as in Figure 1, shown for increasing UV dosage on cells. (Porter: did you divide by the purified DNA signal?) (b) Average TMP signal without torsion from the same genes. (c) Average TMP signal of naked genomic DNA from the same genes. 250 ng of sheared naked genomic DNA was incubated for 2 min with the indicated concentration of psoralen and exposed to UV light for 10 s. The density of adjacent AT or TA dinucleotide cross-linking sites is plotted for comparison with the naked DNA profiles (XL site density, dark blue). (d-f) Percent of 250 ng of input DNA remaining after two rounds of denaturation and treatment with ExoI for each respective condition in (a-c). The response in % DNA remaining is near linear with UV dosage. At low concentrations, we see that the % DNA remaining approaches the background (d-f) as does the signal from the library prep (a-c). This indicates that if the dose is too low, the signal-to-noise limit is being approached. Upon high UV exposure, the % DNA remaining indicates that there are likely multiple crosslinks/fragments (d,e), which may lead to artifacts due to the loss of small fragments during library prep. Upon a Goldilocks dose of UV exposure or psoralen concentration, signals are very similar to each other, indicating that most fragments have exactly one crosslink after Exo digestion (Fig. Supplement 3).

**Figure Supplement 3.**
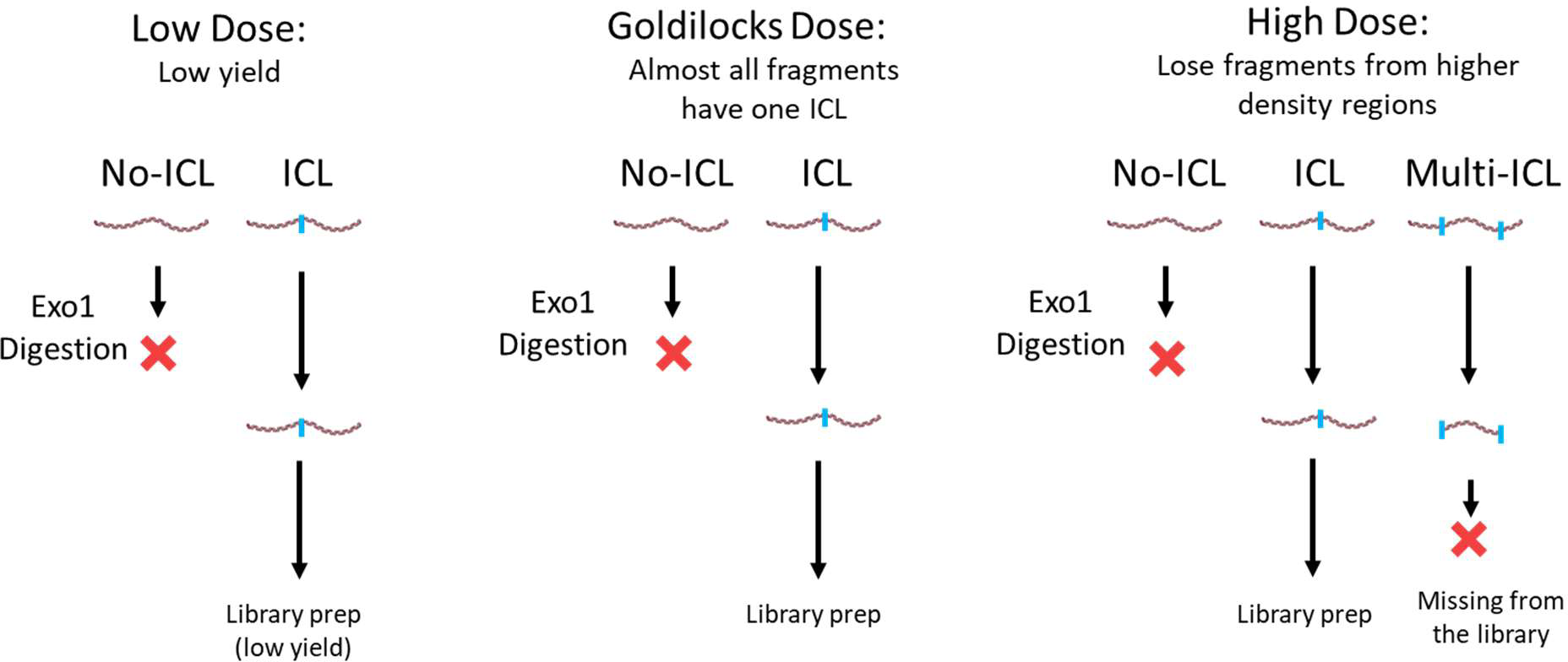
Optimal TMP crosslinking density. Cartoon depicting the optimal inter-strand cross-linking density is when each fragment of DNA has only one TMP cross-link after Exonuclease I digestion.

